# Enhanced delivery of protein therapeutics with a diphtheria toxin-like platform that evades pre-existing neutralizing immunity

**DOI:** 10.1101/2024.04.02.587727

**Authors:** Shivneet K. Gill, Seiji N. Sugiman-Marangos, Greg L. Beilhartz, Elizabeth Mei, Mikko Taipale, Roman A. Melnyk

## Abstract

Targeted intracellular delivery of therapeutic peptides and proteins remains an important but unresolved goal in biotechnology. A promising approach is to engineer bacterial exotoxins that deliver their cytotoxic enzymes into cells and can be engineered to target cancer cells as is the case with immunotoxins. The well-studied diphtheria toxin translocation domain is ideally suited as a delivery platform as it has been shown to be capable of delivering a wide range of macromolecular cargo. Widespread deployment of DT-based therapeutics in humans, however, is complicated by the prevalence of pre-existing anti-DT antibodies from childhood vaccinations that reduce the exposure, efficacy and safety of this important class of protein drugs. Thus, there is a great need for delivery platforms with no pre-existing immunity in humans. Here, we describe the discovery and characterization of a distant diphtheria toxin homolog from the ancient reptile pathogen *Austwickia chelonae* that we have named Chelona Toxin (CT). We show that CT is comparable to DT structure and function in all respects except that it is not recognized by pre-existing anti-DT antibodies present in human sera. Moreover, we demonstrate that the CT translocase is superior to the DT translocase at delivering therapeutic protein cargo into target cells. These findings highlight CT as a potentially class-enabling new chassis for developing safer and more efficacious immunotoxins and intracellular protein delivery platforms for cancer therapy.

## INTRODUCTION

Antibody drug conjugates (ADCs) have emerged as one of the fastest growing classes of targeted cancer therapies, with over a dozen FDA approvals in the past decade^1^. Immunotoxins are a subclass of ADCs with a storied history that dates back to Paul Ehrlich’s original “magic bullet” hypothesis^2–5^. Immunotoxins act by targeting cancer receptors via a receptor binding domain but, in contrast to more traditional ADCs which release a toxic small molecule into cells, immunotoxins deliver a cytotoxic protein into cells. Immunotoxins leverage the architecture of bacterial exotoxins which contain a receptor binding domain (R) capable of binding host cells^6^, a translocation domain (T) that forms a pore in endosomal membranes upon receptor binding and internalization of the toxin^7^, and a cytotoxic domain (C) encoding a highly processive enzyme that is delivered to the cytosol to target and inactivate an important intracellular protein to cause cell death^8^ (**Fig. 1a).**

**Fig. 1.**
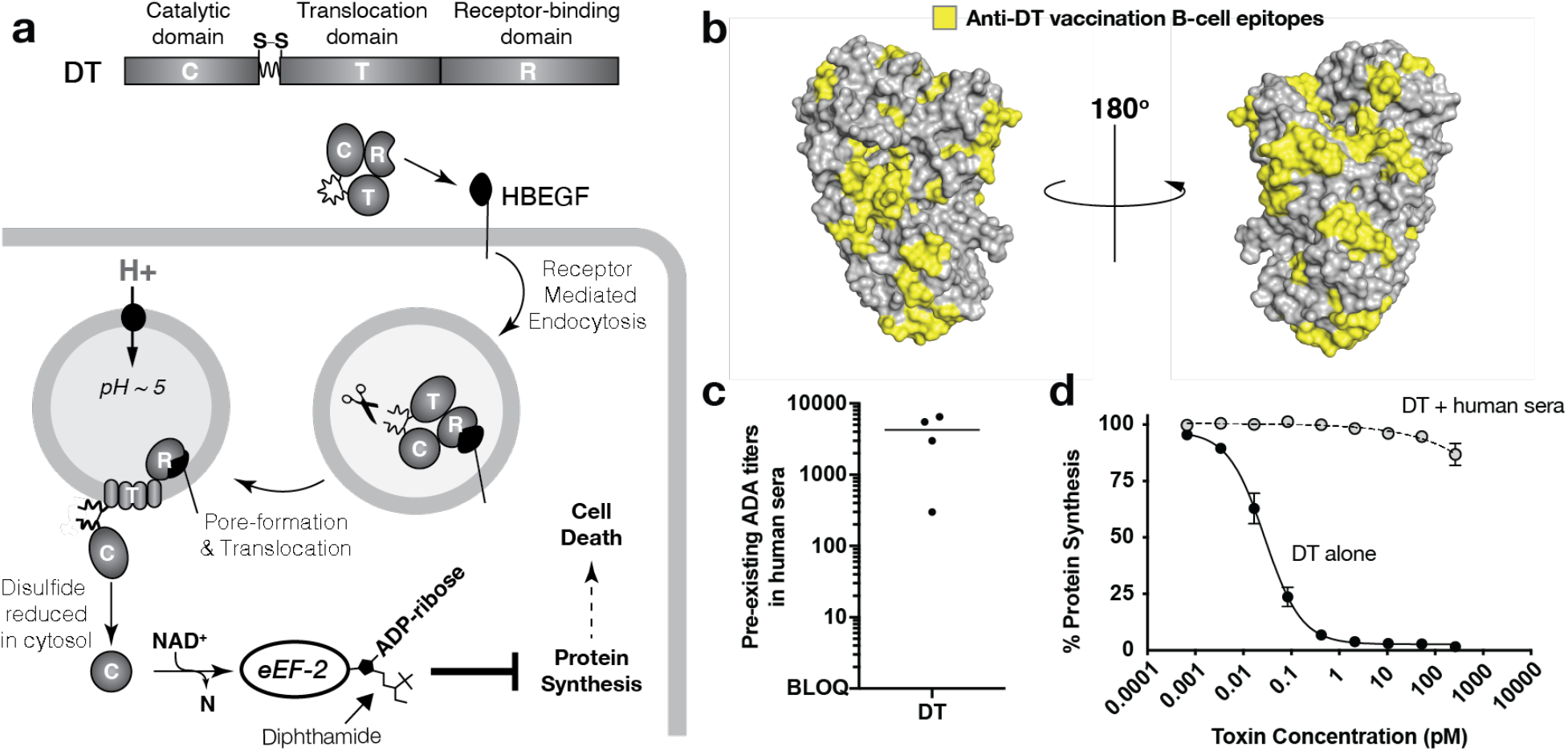
**a,** Molecular mechanism of diphtheria toxin (DT) intoxication pathway. DTs R-domain (DTR) binds HBEGF and undergoes HBEGF-mediated internalization. Furin protease cleaves at a site between the C- and T-domains, and as the endosome acidifies, the DT translocase (DTT) unfolds and inserts into the endosomal membrane to form a pore, through which the catalytic domain (DTC) translocates through and enters the cytosol. Disulfide reduction releases DTC into the cytosol, where it ADP-ribosylates elongation factor 2 (EF-2), inhibiting its function in mediating ribosomal protein translation, thus shutting down protein synthesis in the cell to cause cell death. **b,** B-cell epitopes on DT (yellow) because of vaccination, as characterized by De-Simone et al. **c,** Assessment of IgG binding to DT from pooled human sera, by ELISA. **d,** Human sera neutralization of DT, on cell-based assay. Sera is neutralizing beyond 100pM.

To date there have been three FDA approved immunotoxins: denileukin diftitox, (Ontak™)^9^; tagraxofusp, (Elzonris™)^10^; and moxetumomab pasudotox, (Lumoxiti™)^11^ of which the first two are derivatives of diphtheria toxin (DT). DT is a highly toxic protein secreted by *Corynebacterium diphtheriae*, and the etiologic agent of the disease Diphtheria^12^. Denileukin diftitox and tagraxofusp contain the catalytic and translocation domains of DT, but the receptor binding domain has been replaced with either interleukin-2 (IL-2) or interleukin-3 (IL-3) respectively, which successfully re-targets DT to kill cells expressing their receptors CD25 (IL2RA) or CD123 (IL3RA). As such, Ontak™ was approved for cutaneous T cell lymphoma (CTCL), and Elzonris™ was approved for blastic plasmacytoid dendritic cell neoplasm (BPDCN), increasing survival rates by over 50% for BPDCN patients. Elzonris™ is currently being explored for patients with acute myeloid leukemia (AML)^13^, as AML cells also express high levels of CD123^14^.

To stop the devastating spread of diphtheria in the 20^th^ century, vaccines against DT were developed and are a mainstay in global vaccination programs, resulting in >85% of the global population being vaccinated against DT^15^. DT immunity is achieved by injecting de-activated full-length non-toxic forms of DT (toxoid) into humans, resulting in the development of antibodies that recognize and neutralize DT. In a recent study, the B-cell epitopes on DT were mapped and found to consist of regions in all three domains that cover much of the surface of DT (**Fig. 1b**)^16^. As a demonstration of the natural immunity against DT in a general population, when purified DT is used as bait in an ELISA, we observe high levels of anti-DT titers in pooled human sera (**Fig. 1c**), which neutralize DT-mediated cytotoxicity by over six orders of magnitude (**Fig. 1d**). The neutralizing immunity that protects humans from DT also prevents the widespread use of DT-based therapeutics. Indeed, clinical trial data for both Ontak™ and Elzonris™ highlight the negative impact of pre-existing anti-drug antibodies (ADAs) on an otherwise highly effective targeted therapeutic^9,10^. In total, 66% of CTCL patients had baseline ADAs to Ontak™ of which 45% were neutralizing^9^, and up to 95% of BPDCN patients had baseline ADAs to Elzonris™ prior to treatment^17^. Strikingly, there was a clear inverse relationship between pre-existing ADA titers and drug exposure; in patients with the highest pre-existing anti-DT titers, drug exposure was up to 100-fold lower than in patients with low titers^18^. Moreover, patients with pre-existing ADAs had a lower response rate. Devising strategies to evade pre-existing ADAs would improve all aspects of DT-based therapeutics from pharmacokinetics to pharmacodynamics, ultimately resulting in a better, more durable outcome for cancer patients^19^.

In this study, we screened distant homologs of DT with the goal of identifying a novel toxin-based platform that retained the translocase and catalytic functionality of DT but which was not recognized by pre-existing ADAs in human sera. Two related toxins were identified that are implicated in diphtheria-like diseases in reptiles. We thus investigated this new toxin family as a novel chassis for targeted cancer therapeutics and protein delivery.

## RESULTS

### Functional screening and characterization of diphtheria toxin homologs

To uncover distant relatives with DT-like functionality, we probed the NCBI non-redundant protein sequence database by performing a PSI-BLAST using the 535 amino acid sequence of diphtheria toxin as a query. This search revealed a handful of DT-like putative gene sequences all sharing ∼20-40% sequence identity to DT (**Supplementary Fig. 1; Fig. 2a**). Homologs possessing a predicted translocation domain and a catalytic domain were selected for characterization. To determine the functionality of the individual domains from the DT homologs, we used a “host-guest” platform, wherein the native translocase or catalytic domains from DT (*viz*. the host) were replaced with the corresponding domains from a given homolog (*viz*. the guest) (**Fig. 2b-c**). All DT-homolog chimeras retained the native DT receptor binding moiety (*i.e.,* DT_R_) to normalize the receptor-binding step in the intoxication pathway (**Fig. 1a**). Eight chimeric toxins were expressed and purified to homogeneity in *E. coli*. As a universal readout of function, we quantified inhibition of protein synthesis by the delivered cytotoxic ADP-RT in Vero cells engineered to constitutively express a destabilized luciferase (Vero-NLucP). From eight homologs tested, only two translocases, both from the *Austwickia* genus, were able to deliver DT_C_ at sufficient levels in this context, using minimal linkers and relying on the predicted domain boundaries of homologs (**Fig. 2b**). The two functional homologs are derived from *Austwickia chelonae* and *Austwickia chelonae LK16-18.* Notably, these strains have been reported to cause skin lesions in various reptiles (bearded dragon^20^, king cobra^21^, alligators, turtles^22^, and tortoises), and specifically the strain LK16-18 has been reported to cause cutaneous granuloma in crocodile lizards^23^.

**Fig. 2.**
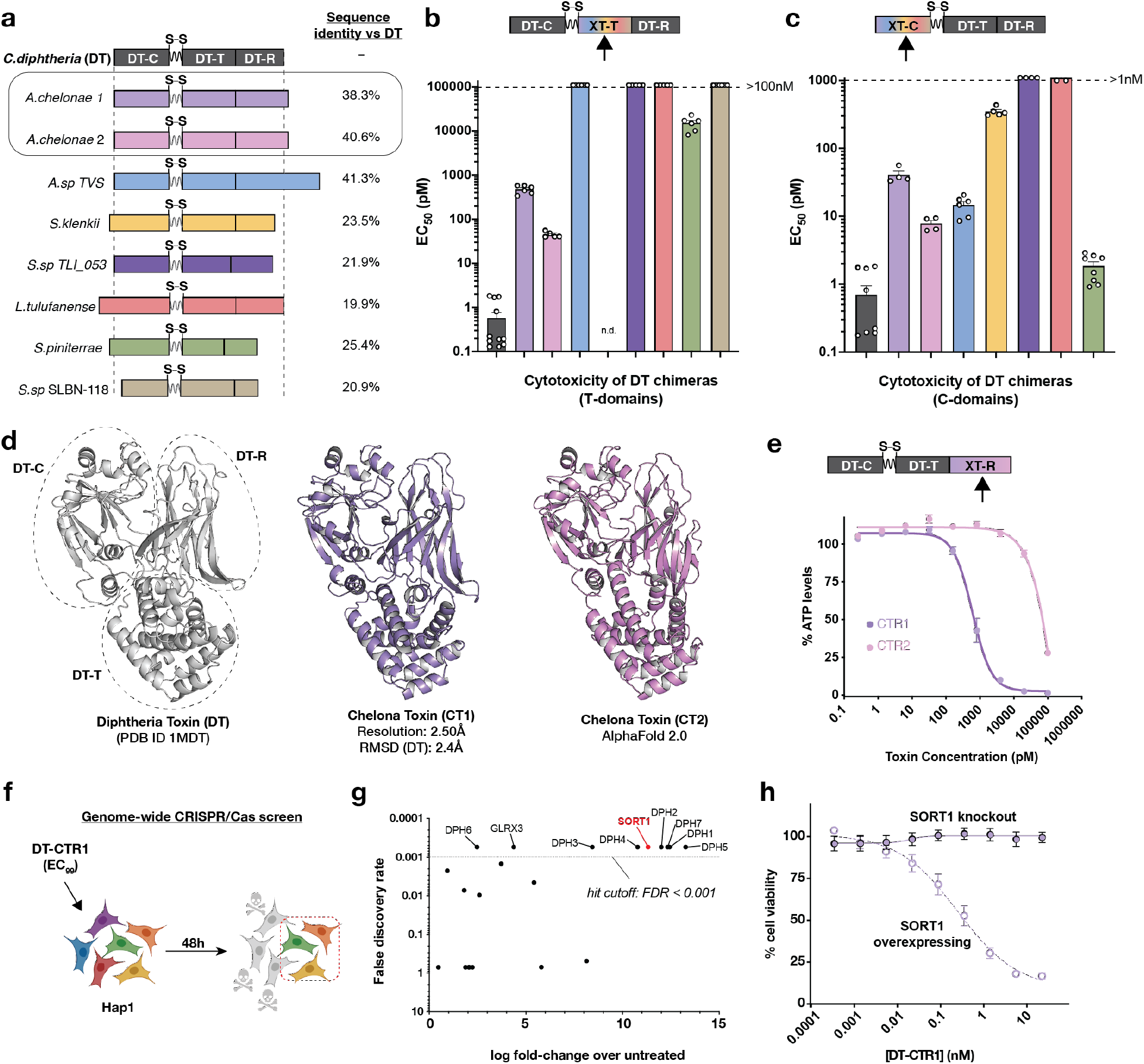
**a,** Sequence identity of putative DT-homologs to DT. Species from which DT-homologs were identified are indicated on the left. Approximate length of each sequence is depicted by the length of the bars, with DT representing 535 residues. **b, c** Screen for functional translocation domains and catalytic domains, respectively. Each chimeric protein was tested on vero-nLucP cells, and EC50 values were plotted on the y-axis. Colours correspond to the respective DT-homolog in **a**. n=3, SEM. **d,** Crystal structure of CT1 and AlphaFold 2.0 structure of CT2. **e,** Testing CT1R and CT2R functionality on human cells. Recombinant proteins were tested on Hap1 cells. CT1R had an EC99 of ∼75nM. **f,** Schematic of the genome-wide CRISPR/Cas9 screen. **g,** Results of CRISPR/Cas9 screen. **h,** Validation of SORT1 as a receptor for CT1R, using Hap1-SORT1 knockout cells and SORT1 overexpressing cells. n=3, SEM.

A similar screen was conducted to identify functional catalytic domains; the catalytic domain of each homolog was recombinantly linked to the translocation and receptor binding domains of DT, and these chimeric proteins were tested on Vero-NLucP cells (**Fig. 2c**). In this case, we found that four of the seven homologs, including those from *Austwickia* were functional to varying degrees. Given their dual functionality, we focused on the DT homologs from *Austwickia*, herein referring to the protein from *A. chelonae* as Chelona Toxin 1 (CT1) and *A. chelonae LK16-18* as Chelona Toxin 2 (CT2).

### Structural characterization of CT family proteins

To better characterize CT1 and CT2, we next sought to elucidate their three-dimensional structures. For CT1, we used hanging drop vapour diffusion and successfully crystallized and obtained 2.50 Å diffraction data. We were unable to use the full-length DT structure (PDB 1MDT) as a search model for molecular replacement; however, using partial search models with 1MDT, we were able to solve the structure (**Fig. 2d**). CT1 has a root square mean deviation (RMSD) of 2.3 Å to DT (401/535 residues aligned) and retains the same three domain, Y-shaped architecture as DT.

For CT2, we used Alphafold 2.0^24,25^ to predict its three-dimensional structure (**Fig. 2d, Supplementary Fig.2**). The RMSD of CT2 to DT was 1.3 Å (383/535 residues aligned) and retains the same Y-shaped architecture as DT.

Both catalytic domains conserve the split β-sheet that is typical of ADP-ribosyl-transferases. The specific residues that have been demonstrated to be important for the catalytic activity of DT_C_ for binding and ADP-ribosylating elongation factor-2 are also conserved in both CT1 and CT2. In comparison to the catalytic domain, much less is known about which residues in the translocation domain are essential for function. That the translocases of CT1 and CT2 share little sequence identity with DT, but retain function, provides an unprecedented opportunity to uncover new functional determinants of the translocase. The translocase is discussed in greater detail later in this study (*vide infra*).

Given the reduced toxicity observed for CT1 and CT2 on Vero cells (**Supplementary Fig.3**) which express high levels of HBEGF, we hypothesized that their receptor-binding domains likely did not recognize the DT receptor. Indeed, the residues implicated in binding to HB-EGF are poorly conserved in CT1 and CT2 (**Supplementary Fig.4**). To determine whether CT1_R_ and CT2_R_ bind a different cell-surface receptor on human cells, we conducted a genome-wide CRISPR/Cas9 screen on Hap1 cells with the TKOv3 library. Chimeras were made in which the R-domains of CT1 and CT2 were recombinantly linked to the catalytic and translocation domains of DT and tested on Hap1 cells (**Fig. 2e**). As a control, we screened wildtype DT (**Supplementary Fig. 5**). As expected, we identified HB-EGF as the top hit along with genes implicated in the biosynthesis of diphthamide – the molecular target for ADP-ribosylation by DT_C_. The CT1 screen also identified diphthamide synthesis genes but did not recover HB-EGF. Instead, among the top hits in this screen was sortilin (SORT1) – a type I membrane glycoprotein trafficking receptor^26^. SORT1 knockout cells were generated and found to be protected from DT(CT1_R_) while remaining susceptible to DT (**Fig. 2h**). Similarly, SORT1 overexpression through lentiviral transduction re-sensitized these cells to DT(CT1_R_) (**Fig. 2h**). These data demonstrate that SORT1 is essential for CT1 toxicity, likely serving as a cell-surface receptor to mediate CT1 entry into cells.

### CT is not recognized by pre-existing anti-DT antibodies in human sera

The structural and functional analysis above shows that CT toxins satisfy the basic criteria set out in this study to identify distant DT homologs having functional cytotoxic enzymes and translocases for developing novel functional toxin scaffolds for therapeutics. Next, we set out to determine the extent to which CT was recognized by pre-existing anti-DT antibodies. Comparison of the known B-cell epitopes on DT to equivalent positions on both CT1 and CT2 shows only one epitope is conserved and 50% of the individual DT epitope residues are different on both CT1 and CT2 (**Fig. 3a, Supplementary Fig.6**). To experimentally determine the degree to which CT-based constructs are recognized by pre-existing anti DT ADAs, we measured the pre-existing ADA titers in pooled human sera against the catalytic and translocation domains in DT (i.e., DT_1-389_) with an equivalent CT-based immunotoxin (i.e., CT_1-389_) (**Fig. 3b**). DT- and CT were immobilized on plates and incubated with human sera, after which an anti-IgG antibody conjugated to HRP was used to determine levels of bound ADAs. Whereas the DT-based scaffold showed titers of ∼10^6^, the titers against the equivalent CT-based scaffold were below the limit of quantification.

**Fig. 3.**
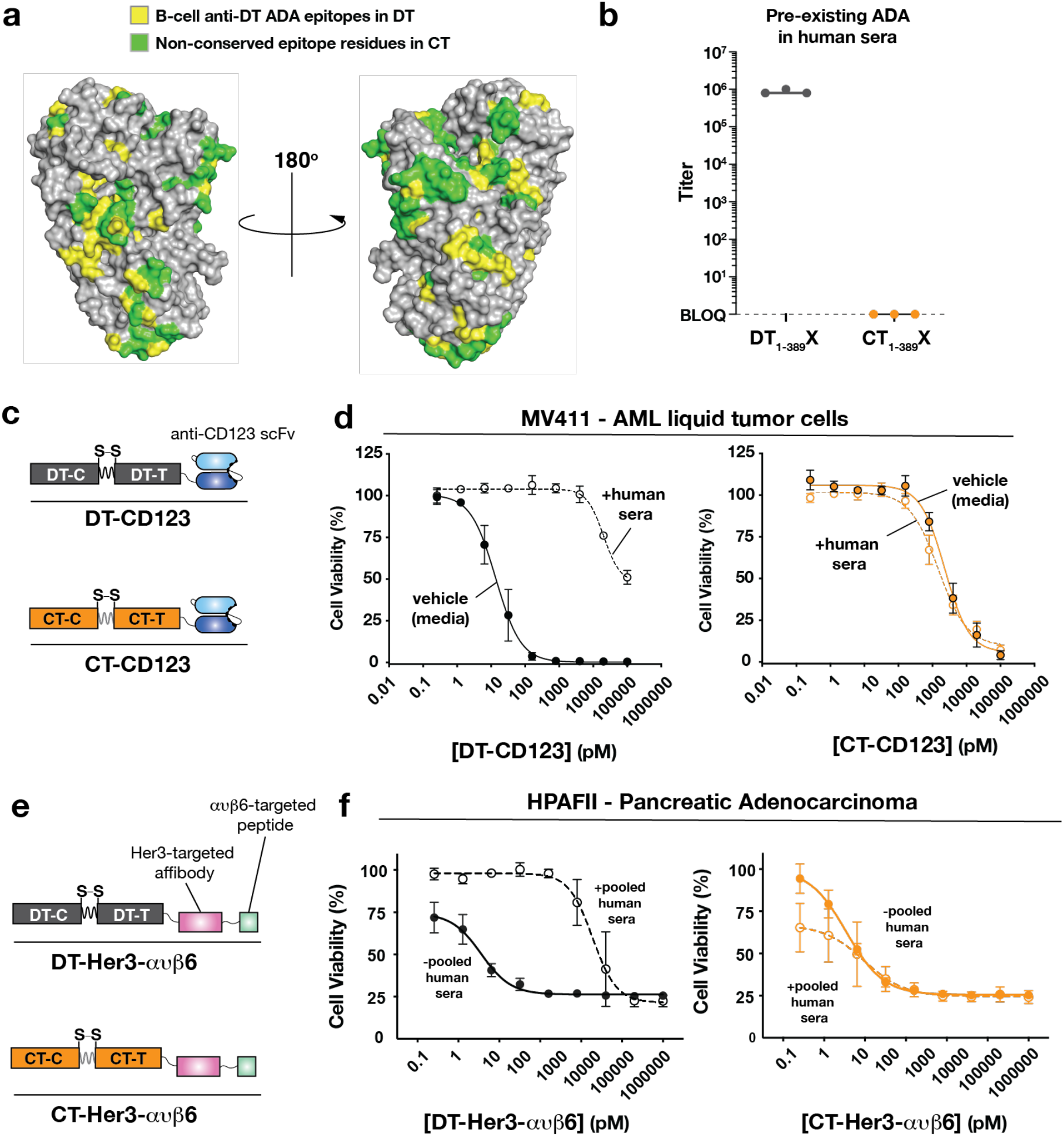
**a,** Conservation of epitopes between DT and CT1. DT (pdb 1mdt) is used as the backbone. In yellow are epitopes conserved in CT1, and in green are epitopes that are not conserved. **b,** ELISA showing binding of human sera to either the catalytic domain and translocation domain of DT or CT. DT1-389 is the first 389 residues of DT which contain the C- and T-domains of DT. CT1-389 is the catalytic domain of *A.sp TVS* and translocation domain of CT2. X in both cases is the scFv binding CD123. **c,** Diagram of the DT- and CT-based immunotoxins binding CD123. **d**, CD123 binding immunotoxins tested on AML cells, in the presence and absence of human sera. DT is significantly neutralized in the presence of human sera, whereas CT is not. n=3, SEM. **e,** Diagram of DT- and CT-based immunotoxins binding HER3 and integrin ανβ6. **f**, Immunotoxins tested on pancreatic cancer cells, in the presence and absence of human sera. DT is significantly neutralized in the presence of human sera, whereas CT is not. n=3, SEM.

### CT-based immunotoxins are functional and evade neutralization by pre-existing ADAs

While it was important to demonstrate that CT was not recognized by pre-existing ADAs in human sera, it was similarly important to have a modular protein delivery chassis that could accommodate various receptor binding domains. We thus constructed two different retargeted DT- and CT-based immunotoxins for liquid and solid tumors. CD123 – the target of Elzonris™ – was used for liquid tumors^27^ (**Fig. 3c**). DT-CD123 and CT-CD123 were tested on MV-4-11 AML cells in the absence and presence of human sera, and cell viability was measured. While both immunotoxins were toxic to cells, DT-CD123 was potently neutralized by sera while the CT-CD123 was not (**Fig. 3d**). The solid tumour receptors HER3^28^ and integrin ανβ6^29^ were also similarly targeted, on pancreatic cancer cells (**Fig. 3e**). As above, in the presence of human sera, the DT-based immunotoxin was significantly neutralized but the CT-based immunotoxin was not (**Fig. 3f**). Taken together, these experiments demonstrate that unlike DT-based immunotoxins, ADAs in human sera do not neutralize or bind CT-based immunotoxins, supporting CT as an alternative immunotoxin platform to DT. Importantly, the finding that DT constructs lacking DT-R were potently neutralized highlights the fact that ADA binding to the catalytic and/or translocation domains of toxins are sufficient for neutralization.

### Exploring the potential of CTT as a next-generation immunotoxin chassis

In the traditional immunotoxin design, the therapeutic payload that gets delivered into the cytosol of the target cell is the native cytotoxic enzyme of the toxin, which inhibits protein synthesis. Recently, groups including ours, have demonstrated the potential of DT_T_ to deliver other protein cargo of interest into cells, expanding the therapeutic utility of DT^30–33^. To evaluate the ability of CT_T_ to serve as a general translocase of diverse protein cargo of therapeutic interest, we designed DT and CT-based translocase constructs flanked by flexible linkers that were retargeted with either a HER3-targeting affibody^28^ or a TEM8-targeting scFv^34^ **(Fig. 4a, b**). Four differently-sized protein cargoes were selected, including DT_C_ itself; RRSP, an endopeptidase that cleaves all Ras isoforms and Rap1; and the glucosyltransferase domains (GTDs) from *C. difficile* Toxin B (TcdB-GTD), and *C. perfringens* large toxin (TpeL-GTD) that target different intracellular small GTPases – many of which are implicated in different malignancies. Each cargo was placed to the amino terminus of a cassette consisting of a flexible linker and the furin site flanked by cysteines that form a disulfide bond from DT (**Fig. 4a**). The four HER3 targeted chimeras were tested on human pancreatic HPAF-II cells (high HER3 expression), and the four TEM8-targeted chimeras were tested on human neuroblastoma SKNAS cells (high TEM8 expression). After a 72-hour treatment, the cell viability of each of the chimera pairs was compared (**Fig. 4c).** Remarkably, in all cases, CT_T_ outperformed DT_T_ in delivering a particular protein cargo, showing increased potency on all cells tested, indicating that CT_T_ is a more efficient protein translocase than DT_T_.

**Fig. 4.**
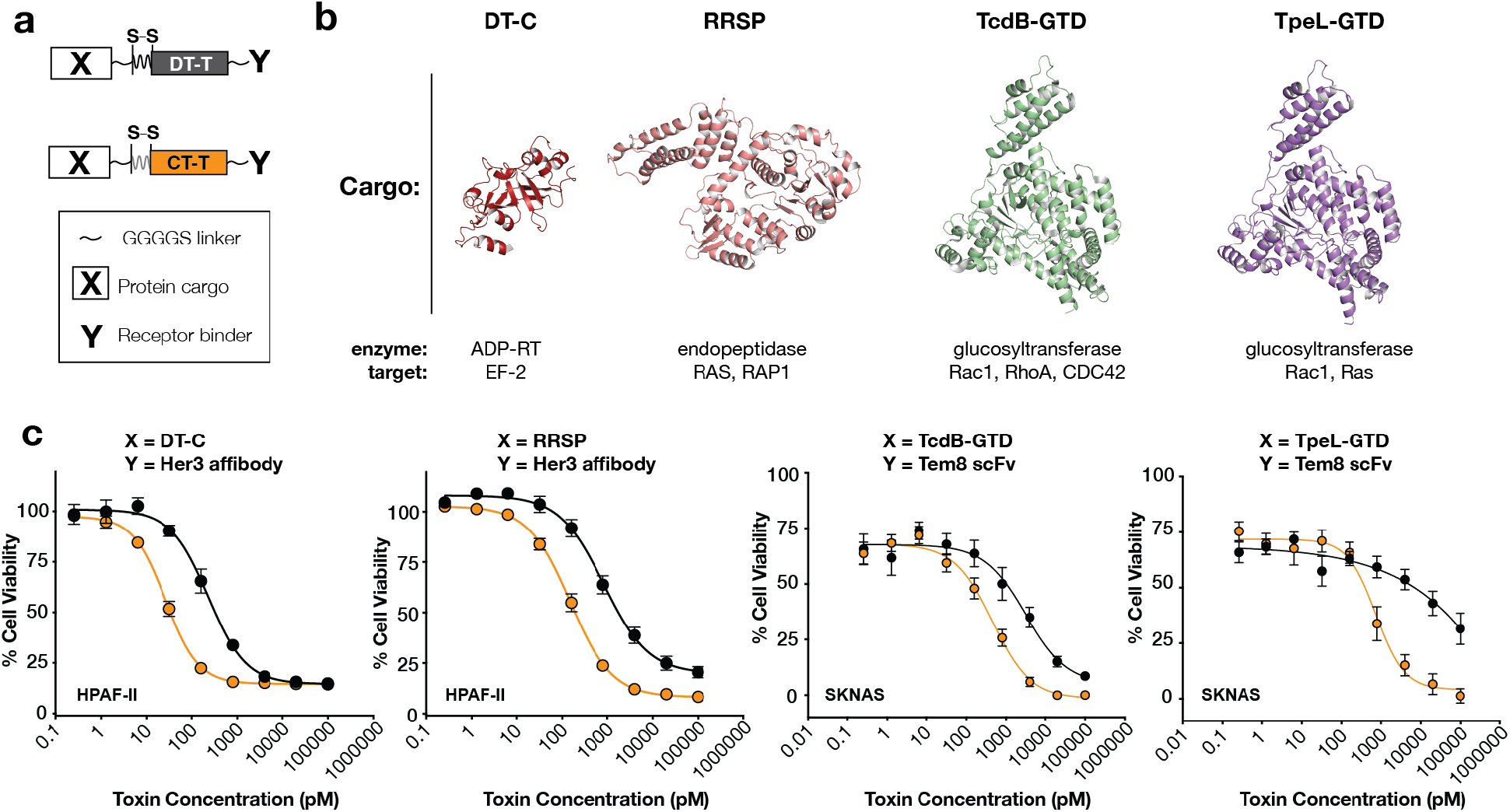
**a,** Schematic of designed immunotoxins delivering diverse cargo. **b,** Structures and functions of catalytic domains (cargo) delivered by either the DT or CT translocases. PDB IDs for the structures are as follow: DTC: 1mdt, RRSP: 5w6l, TcdB-GTD: 7s0z, TpeL-GTD: **XXXX**. **c,** Cellular delivery of each cargo by the translocation domain of DT or CT, using various receptor binding domains. n=3, SEM.

### The CT translocase enables “earlier” endosomal escape of cargo

To elucidate the basis for the enhanced delivery by CT_T_, we further evaluated the structure and function of the two CT translocases. Similar to what is seen in DT_T_, CT translocases maintain the nine α-helices that are organized into three distinct layers (**Fig. 5a**). Models for the pH-dependent unfolding and insertion of the translocation domain have been proposed, and the roles of aspartic acid 352, glutamic acid 349, and six histidines (residues 223, 251, 257, 322, 323, 372) have been elucidated^35–37^. Glu-349^38^ and Asp-352^39^ are thought to be important in driving and maintaining the insertion of the translocase in the endosomal membrane, and both are conserved in CT1 and CT2 (**Fig. 5b**). Interestingly, however, only half the histidine residues are conserved in the CT translocases (**Supplementary Fig.7**). It was previously reported that the unfolding of DT_T_ at a pH <6.0 is mediated by the local effects of His-223 (the “safety latch”) that lowers the pKa of His-257^35^. In both CT1 and CT2, the safety latch is absent; in the position occupied by a histidine in DT, there is a phenylalanine in both CT variants (**Fig. 5c**). It was shown previously that disrupting the safety latch in DT results in an increase in the pH at which DT_T_ unfolds^35^. To assess whether CT unfolds at a higher pH than DT, we used differential scanning fluorimetry (DSF)^40^ and assessed the melting temperature of DT or CT at pH 5.0–7.5. We observed a shift in the pH-dependent unfolding profile of CT relative to DT, suggesting that CT initiates the unfolding process at a higher pH than DT (**Fig. 5d**).

**Fig. 5.**
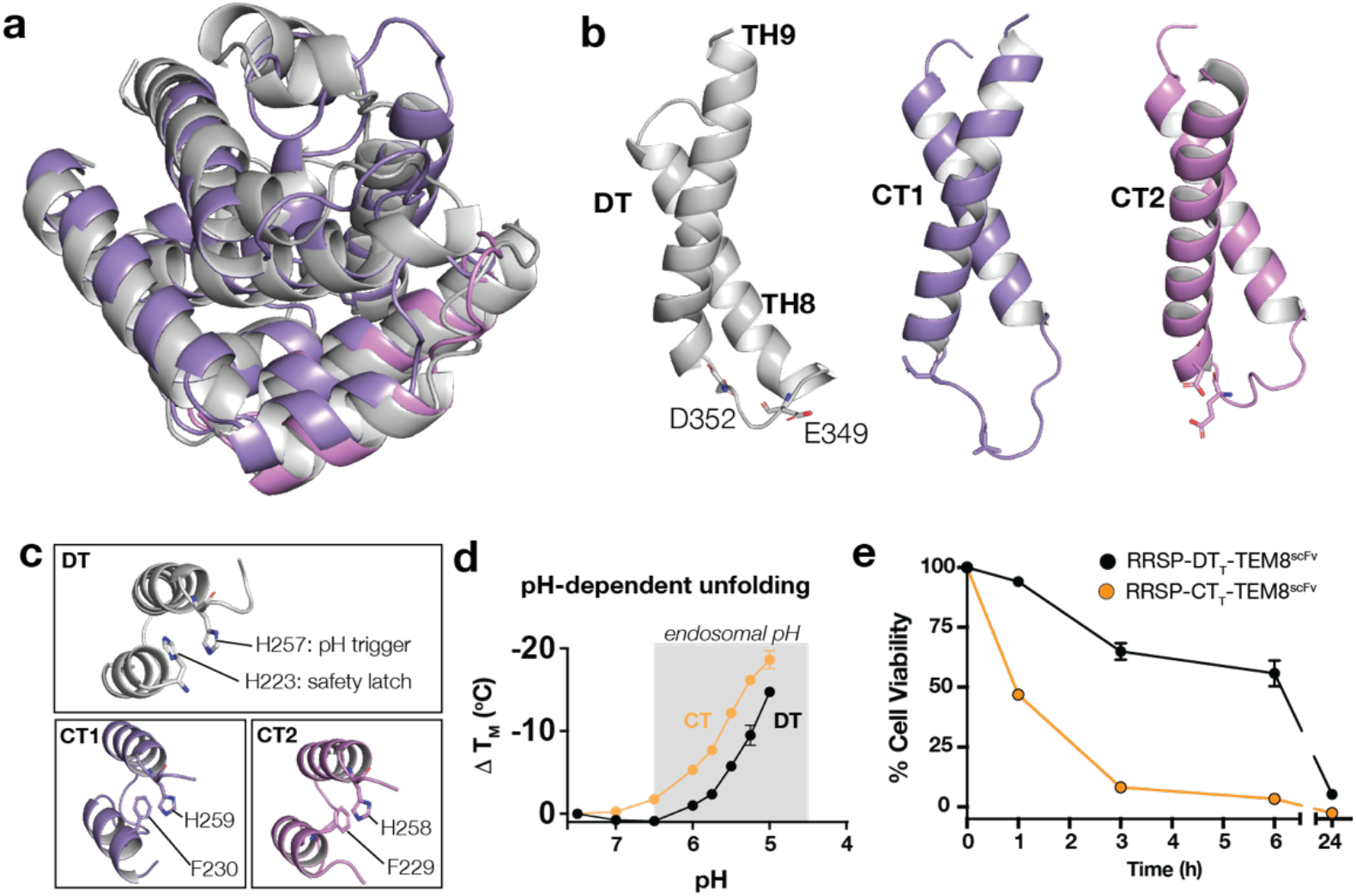
**a,** Structure overlay of translocases from DT (grey), CT1 (purple) and CT2 (pink). **b–c,** Key functional residues are depicted in stick format. Colours correspond to a. **d,** Melting temperature of DT and CT1, by differential scanning fluorimetry. **e,** Kinetics experiment on HCT116 cells. Time points indicate how long toxins were incubated with cells before replacing cells with toxin-free media. ATP levels were measured at 72hrs and plotted as cell viability on the y-axis.

We hypothesized that the higher pH of unfolding for CT-based translocases may enable earlier, or more rapid escape from endosomes that become increasingly acidic as they transit from the cell surface toward late endosomes and lysosomes. To assess if the kinetics of translocase delivery differed, DT_T_- or CT_T_-based immunotoxins were incubated with cells for various timepoints, after which the proteins were washed off and cells were left to grow for 72 hours before reading for viability. CT_T_ delivered RRSP rapidly into cells, reducing the 72-hour viability of HCT116 cells by ∼50% after just one-hour incubation versus DT_T_, which reduced the 72-hour cell viability by ∼40% after 6 hours of incubation (**Fig. 5e**).

## DISCUSSION

The work presented herein arose from our desire to circumvent the liability of pre-existing ADAs to current immunotoxin platforms. Our approach to do a functional screen of DT homologs uncovered toxins from the *Austwickia* genus as promising candidates to replace DT as a potential alternative immunotoxin platform. These novel immunotoxins are not neutralized or recognized by ADAs in human sera and retain equivalent or superior functionality to DT in their capacity as immunotoxins and protein delivery vectors. While *Austwickia chelonae* is a reptile pathogen, it has never been reported as a human pathogen and our data suggest that humans likely have not been exposed to these toxins. This is unlike exotoxin A, the other major toxin used in immunotoxins. Despite not being immunized against exotoxin A, many humans harbour pre-existing ADAs due to prior exposure to *Pseudomonas aeruginosa*, a commensal pathogen in humans^11^.

Solid tumours remain one of the hardest classes of cancer to target, and immunotoxins targeting solid tumours suffer substantially from ADAs^41^. In this study we targeted HER3, integrin ανβ6, and TEM8 expressing cancer cells, highlighting the versatility of CT-based immunotoxins to retargeting. An intriguing finding of this study was that CT can use SORT1 as a cell-entry receptor for intoxication. Fortuitously, SORT1 is being explored as a target for ovarian cancer^42^. SORT1 is also being explored as a receptor for protein and siRNA delivery, using galectin-1^43^. Improving the binding of CT_R_ to SORT1 through affinity maturation approaches may yield a novel immunotoxin with all three native domains of CT being used.

While this study addresses the issue of pre-existing ADAs to immunotoxins, the development of ADAs to CT during treatment is expected, as CT is of bacterial origin. To limit ADA development to DT-based therapeutics, many groups have attempted to remove B cell epitopes on DT – in one such study, seven mutations were made to DT that made it less immunogenic while retaining most of its activity^44^. In CT1 and CT2, six of these seven residues are not conserved, and so perhaps CT1 and CT2 are not as immunogenic as DT. Many groups are also investigating strategies to delay the onset of ADA production by co-dosing with immunosuppressive drugs^45^. Future studies will explore the *in vivo* immunogenicity of the CT immunotoxin platform.

Another issue often faced by immunotoxins (and other therapies such as IL-2 cytokine therapy) is vascular leak syndrome (VLS). Baluna et al. conducted a study in which they used ricin A chain and IL-2 cytokine, both of which cause VLS, to identify a consensus sequence (*x*D*y*, where *x* could be the amino acids Leu, Ile, Gly, or Val, and *y* could be Val, Leu, or Ser) that was linked to VLS in a cell-cell adhesion assay^46^. The catalytic domain of diphtheria toxin has a N-terminal VDS sequence, and when Cheung et al. made a Val to Ala mutation, they observed an increase in the maximum tolerated dose of Ontak(V6A) vs Ontak™ in a mouse B16F10 melanoma model^47^. Upon a deeper evaluation of the sequence of the catalytic domain from *A.sp TVS* we found that it has no *x*D*y* motifs. We hypothesize that VLS may be less of an issue with the catalytic domain of *A.sp. TVS*, and future studies will explore strategies to limit VLS in patients.

## CONCLUSION

Pre-existing ADAs to immunotoxins limit the pharmacokinetics, efficacy and safety of this powerful class of cancer therapeutics. This study sought to address that liability by searching for an alternative immunotoxin platform that did not suffer from baseline ADAs that current DT- and exoA-based platforms suffer from. We identified bacterial toxins from *Austwickia chelonae* and demonstrated that they conserved all the functional machinery required of an immunotoxin platform. We built Chelona Toxin (CT)-based immunotoxins targeting both liquid and solid tumours and showed that they are not neutralized by ADAs in human sera. Finally, we demonstrated the superior potential of the translocase of CT to deliver various therapeutically interesting cargo into cells, highlighting its capacity as a novel protein delivery chassis.

## MATERIALS and METHODS

All cell lines were purchased from ATCC and maintained as per manufacturer instructions.

### DT homolog screens – construct generation and purification

Accession numbers for all DT homologs in order of Fig. 2a, are as follow: WP_143115263.1/GAB79386.1, WP_116115734.1, WP_219106995.1, WP_120757473.1, SDT83331.1, WP_189053160.1, JZ58907.1, and WP_160159328.1. Of note, *A.chelonae* was originally separated into a DT_A_-like and DT_B_-component, by a one nucleotide insertion (accession number NZ_BAGZ01000024.1 G43277, compliment). This point insertion was removed prior to codon optimization.

Each sequence was aligned to the DT sequence to determine the identity percentage and boundary cut-offs for chimeric proteins shown in Fig. 2b and 2c. BLOSUM 62 was used to calculate percent identity. The DT_T_ boundaries were from residues 202–377 and DT_C_ boundaries were from residues 1–185. Genes were cloned into the Champion™ pET SUMO *E. coli* expression system by Bon Opus Biosciences. A 2mL starter culture was inoculated into 35mL of LB medium and induced with 1mM IPTG at 25C for 4hours. Cells were centrifuged at 5000rpm and resuspended in lysis buffer (1% protease inhibitor cocktail, 1mg/mL lysozyme, 0.01% Pierce™ universal nuclease inhibitor, 10mM imidazole, 300mM NaCl, 20mM Tris-HCl pH 8.0). Cells were lysed by sonication. Whole cell lysate was centrifuged at 18000xg and the supernatant was incubated with Pierce™ Ni-NTA Magnetic Agarose. The protein was eluted with 300mM imidazole, buffer exchanged into 300mM NaCl, 10mM Tris-HCl pH 8.0, and incubated with SUMO protease overnight at 4C, to cleave the 6xHis-SUMO affinity tag. The protein was incubated with Pierce™ Ni-NTA Magnetic Agarose and the flowthrough (protein) was collected.

### DT homolog screens – protein synthesis assay

Vero-nLucP cells were engineered as previously described^40^, and were plated at 5000 cells/well in 96-well white clear bottom plates (Corning). The following day, protein toxin was added and incubated for 24 hours, after which cells were read for luminescence signal using the NanoGlo® Luciferase Assay kit (Promega), on a SpectraMax M5e plate reader (Molecular Devices). Data was corrected to untreated cells (100% nanoluciferase signal). EC_50_ values were determined by Prism software (GraphPad).

### Crystallization and structure determination of CT1

The CT1 gene is from accession numbers WP_143115263.1 and GAB79386.1. The *E. coli* codon optimized gBlock gene fragment was ordered from Integrated DNA Technologies and cloned into the Champion™ pET SUMO *E. coli* expression system by Gibson Assembly.

A 50mL starter culture of NiCo21 (DE3) *E. coli* cells (New England Biolabs) was inoculated into 1L of LB medium and induced with 0.1mM IPTG at 18C for 18hours. Cells were centrifuged at 5000rpm and resuspended in lysis buffer (1% protease inhibitor cocktail, 1mg/mL lysozyme, 0.01% Pierce™ universal nuclease inhibitor, 20mM imidazole, 500mM NaCl, 20mM Tris-HCl pH 7.5). Cells were lysed with three passes through an Emulsiflex C3 (Avestin) at 15000 psi. Whole cell lysate was centrifuged at 18000xg and the supernatant was filtered through a 0.45um filter and passed over a HisTrap FF crude column (Cytiva). The protein was eluted with 50-75mM imidazole, buffer exchanged into 150mM NaCl, 20mM Tris-HCl pH 7.5, and incubated with SUMO protease overnight at 4C, to cleave the 6xHis-SUMO affinity tag. The protein was flowed over a HisTrap FF crude column and the flowthrough (protein) was collected and concentrated to 8 mg/mL by centrifugation. Data collection and model refinement statistics are in Supplementary Table 1.

Hanging drop vapour difsfusion was used to grow crystals. The condition in which CT1 crystals were obtained contained 2uL of mother liquor (0.2M calcium chloride, 0.1M Tris-HCl pH 8.5, 25% (w/v) PEG4000) and 1uL of 8 mg/mL protein. The drop was dehydrated over 130uL of 2M (NH_4_)_2_PO_4_ for 45 minutes prior to freezing in liquid nitrogen. Data was collected at the Advanced Photon Source on the 23-ID-D beamline at a ^43^ of 1.03319Å.

Initial phases were determined using Phaser in the Phenix software package by using a multi-component search models with individual DT domains (C-domain residues 13-167, R-domain residues 391-531, T-domain residues 205-378) in which disordered loops had been removed. The structure was refined using iterative cycles of phenix.refine and autobuild.

### Structure determination of CT2 using AlphaFold 2.0

The google.colab notebook was used to obtain the structure of CT2^24,25^. Sequence from accession number WP_116115734.1 was used as the query sequence, the the pdb100 was set as the template mode.

### Receptor binding domain cell sensitivity assays

Receptor binding domain boundaries for CT1 and CT2 were determined by aligning each sequence to DT_R_. Cloning was performed by Bon Opus Biosciences into the Champion™ pET SUMO *E. coli* expression system, and proteins were expressed similar to the DT homolog screen. Hap 1 cells were seeded in a 96-well plate one day prior to application of toxin at ∼35-40% confluency. Serial dilutions of individual toxins were made in storage buffer (150 mM NaCl, 20 mM Tris pH 7.5, 5% glycerol) and incubated with cells for 48hr. Toxin sensitivity was measured with CellTiter-Glo reagent (Promega) according to manufacturer protocol. Viability was normalized to untreated wells prior to data visualization with Prism software (GraphPad).

### Virus generation

To generate lentivirus for the construction of the Hap1 TKOv3 library^48^, a ∼1:1:1 molar ratio mixture of library transfer (lentiCRISPRv2; Addgene plasmid #52961) and packaging plasmids (psPAX2, Addgene #12260; pMD2.G, Addgene #12259) was prepared in serum-free media (Opti-MEM™; Gibco™, cat # 31985062). A ∼3:1 ratio of X-tremeGENE™ 9 DNA Transfection Reagent (Roche, XTG9-RO) was added to the mixture prior to incubation and application on HEK293T cells using standard methods ^1^. Virus was collected 48hr after infection. To generate lentivirus for single gene knockout lines, gRNAs were selected from the TKOv3 library and cloned into lentiCRISPRv2 under established protocol ^1^. Packaging and lentiCRISPRv2 plasmids were transfected into HEK293Ts using X-tremeGENE™ 9 as above. Similarly, to generate lentivirus for HEK293T SORT1 overexpression cell lines, SORT1 cDNA was obtained from the hORFeome V8.1 collection and cloned into pLX301 (Addgene #25895) for co-transfection with psPAX2 and pVSV-G (Addgene #138479). Virus was collected from HEK293Ts 72hr post-transfection for all gene-specific constructs.

### Hap1 genome wide CRISPR KO screens

To generate a starting (T0) cell population with ∼200-fold gRNA library coverage, 50 million Hap1s were seeded and transduced the same day with the lentiviral TKOv3 library at MOI 0.3 under 8 μg/mL polybrene. The day after infection, fresh media containing 1 μg/mL puromycin was applied for 48hr to eliminate cells without gRNA inserts. Surviving T0 cells were pooled, and the library was maintained at minimum ∼100-fold coverage beyond this point. Cells underwent expansion before 7 million cells were seeded into two 15 cm plates per condition on T4. Toxin was applied to cells the next day (T5); at T7, toxin-containing media was removed and replaced with complete growth media (IMDM, 10% FBS) to allow survivor repopulation. Untreated cells were passaged every 72hr in parallel with the replacement of fresh media on toxin-treated plates. Survivors were reseeded once colonies were visible and collected when plates reached ∼80% confluency alongside untreated cells from the same day.

### Next-generation sequencing library preparation

Genomic DNA was extracted using the Wizard® Genomic DNA Purification Kit (Promega, cat #A1120) following manufacturer protocol. gRNA inserts were then amplified from each sample. Amplicons were barcoded with Illumina TruSeq adapters i5 and i7 sequences prior to NGS sequencing at a read depth of at least 5 million reads per sample. Hits were identified from FASTQ files and ranked using MAGeCK software^49^.

### Generation of stable cell lines for screen validation

Hap1 or HEK293Ts were seeded in 6-well plates at < 40% confluency. The next day, cells were transduced with construct-specific virus and 8 μg/mL polybrene. After overnight incubation, virus-containing media was removed, and cells were selected with 2 μg/mL puromycin for 48hr. Each polyclonal stable cell population was reseeded in 10 cm plates for expansion. Monoclonal cell lines were obtained from polyclonal populations through serial dilutions in 96-well plates. Clones were validated through immunoblot for target proteins.

### ELISA with human sera

Nunc MaxiSorp™ plates (Thermo Fisher Scientific) were immobilized with 1ug of protein overnight at 4C, in carbonate pH 9.4 buffer. They were then blocked with 3% milk in TBS (TBSM) overnight at 4C. Dilutions of human serum (Human Serum age 4-6, Innovative Research) were performed in TBSM, and wells were incubated with 100uL for two hours at 37C. Of note, various human patient samples were used in the ELISAs, and immunization status of the patients was unknown. Wells were then washed with TBST (0.1% tween 20) and then incubated with an anti-human IgG antibody conjugated to HRP (Abcam, ab102420) for one hour at 37<u>°</u>C, that was developed using 100uL of TMB reagent (Thermo Fisher Scientific). Absorbance was read at 630nm. Background was twice the A630 reading of control wells (no-protein, +human serum; bovine serum albumin, + human serum).

### Serum toxicity assays

Protein toxins were incubated with either human sera (Human Serum age 4-6, Innovative Research) or PBS in a 1:1 ratio, for 30 minutes at room temperature. Sample was then added to cells that had been plated at 10,000 cells/well the previous day, in a 96-well white clear bottom plate (Corning). Cells were incubated for 72 hours, upon which cells were read for viability. Viability of MV-4-11 cells was assessed using the CellTiterGlo 2.0 kit from Promega (as per manufacturer instructions); viability of HPAFII cells was assessed using PrestoBlue reagent from Thermo Fisher (as per manufacturer instructions). Values were corrected to sera-only treated cells, which represented 100% viability.

### CD123 and TEM8 targeting immunotoxin generation

The immunotoxins targeting CD123 use an scFv as the receptor binding domain. The scFv was derived from the antibody used in the ADC IMGN632^27^ – the variable light chain (V_L_) was linked to the variable heavy chain (V_H_) via a (G_4_S)_3_ linker. The V_L_-V_H_ scFv was linked to the bacterial toxin also via a (G_4_S)_3_ linker. DT-CD123 was made using the first 389 residues of DT. CT-CD123 was made using residues 32–217 from CT-TVS, 186–201 from DT, and 204–379 from CT2. The scFv targeting TEM8 was derived from the m825 antibody described in Szot et al. similar to the CD123, except the orientation was V_H_-V_L_. All cloning was performed by Bon Opus Biosciences, into the pET SUMO *E. coli* expression system. Proteins were expressed and purified similar to the DT homolog screen. TEM8 targeting constructs were expressed at 18C for 18hours.

### Differential Scanning Fluorimetry (DSF)

DSF with DT and CT1 was performed as previously described^40^. Briefly, DT or CT1 proteins were diluted in 150mM citrate phosphate buffer containing 5X SYPRO Orange (Invitrogen). The pH of the buffer varied from 4.0–7.5, in 0.5 pH increments. Fluorescence of SYPRO Orange was measured using a BioRad CFX96 qRT-PCR thermocycler, and the BioRad CFX Manager 3.1 software was used to determine melting temperatures. Melting temperature data was normalized to the melting temperature at pH 7.5 for each respective protein.

### Crystal structure of TpeL-GTD

The TpeL-GTD gene was cloned from accession number BAF46125.1 (residues 1–543), into the pET-28a expression plasmid (EMD Millipore). Protein was expressed and purified as described above. The 6xHis tag was not removed. Hanging drop vapour diffusion was used to grow crystals. TpeL-GTD crystallized in 0.2M potassium sodium tartrate and 20% (w/v) PEG 3350 (1uL mother liquor, 1uL of 16mg/mL protein). Crystals were crushed and re-seeding into the mother liquor for optimization. Crystals were frozen in liquid nitrogen. Data was collected at the Advanced Photon Source on the 23-ID-D beamline at a wavelength of 1.0332Å. Initial phases were determined using Phaser in the Phenix software package by molecular replacement with the structure from PDB ID 4DMV (the GTD from *C. difficile* toxin A). The structure was refined using iterative cycles of phenix.refine and autobuild, to a final resolution of 1.9Å. Data collection and model refinement statistics are in Supplementary Table 1.

