## Supplemental Figures for "Enhanced delivery of protein therapeutics with a diphtheria toxin-like platform that evades pre-existing neutralizing immunity"

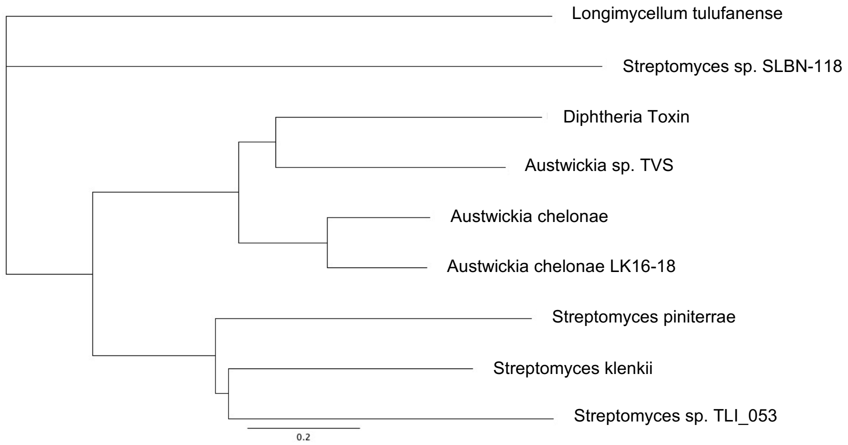


**Supplementary Fig. 1| Phylogenetic tree of DT homologs.** The sequences of each DT homolog were aligned using Blosum62, and a phylogenetic tree was generated using the Geneious 11.0.5 Tree Builder using the Jukes-Cantor genetic distance model and the Neighbour-Joining tree build method. DT homolog proteins are named by the species from which they were extracted. Austwickia genus proteins are the closest to DT.

**a**

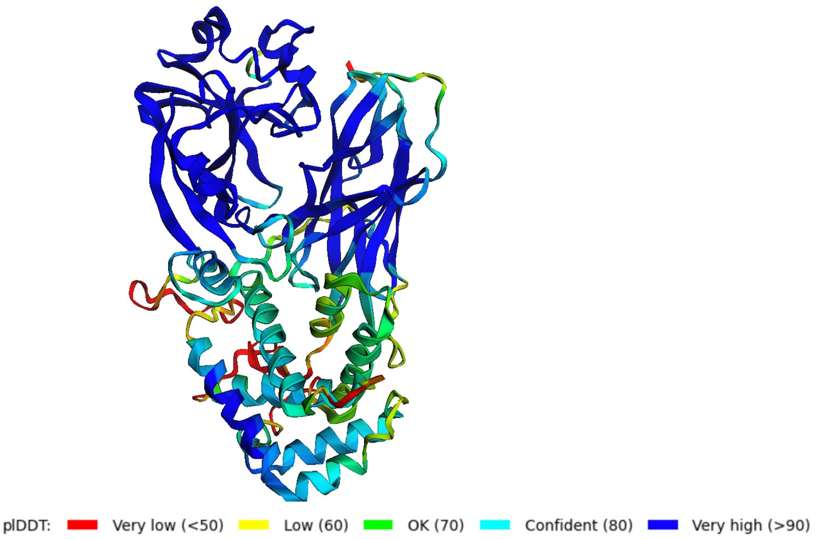


**b**

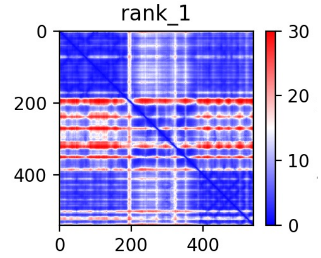


**Supplementary Fig. 2| AlphaFold 2.0 error scores.** The sequence of CT2 was inputted into the online colabfold notebook. a, Predicted local distance difference test (pIDDT) score for rank 1. The translocation domain is predicted the poorest, and the C- and R- domains are predicted with high probability. The corresponding predicted alignment error (PAE) score for rank1, where blue represents low alignment error, and red represents high alignment error. Overall, there is low error in the structure predicted in rank 1.

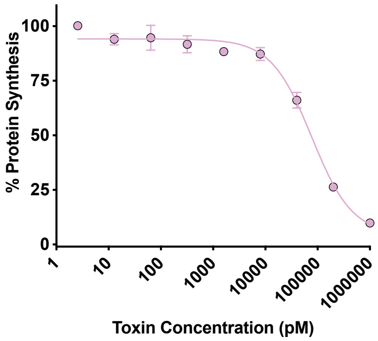


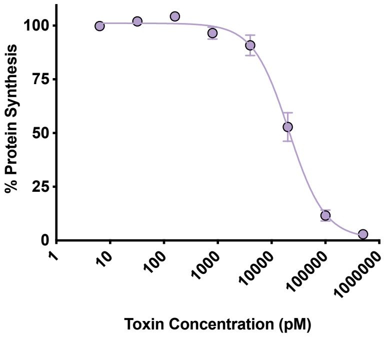


**Supplementary Fig. 3| Toxicity data of CT1 and CT2**. Full length CT1 and CT2 were purified and tested on Vero-nLucP cells, and luminescence signal was measured at 24 hours post treatment. Both CT1 (pink) and CT2 (purple) have $EC_{50}$ values >50nM.

| | 391 | 430 | 433 | 464 | 465 | 468 | 470 | 510 | 512 | 516 | 523 | 526 | | |
|---|---|---|---|---|---|---|---|---|---|---|---|---|---|---|
| | H | A | L | I | D | V | F | G | L | K | V | K | DT | |
| | S | V | A | T | E | L | F | D | L | I | T | K | CT1 | **% identity** = 23.1%; **% similarity** = 45.0% |
| | S | T | M | A | G | L | F | D | L | I | T | K | CT2 | **% identity** = 28.0%; **% similarity** = 47.8% |

**Supplementary Fig. 4| Sequence alignment of key functional residues in receptor binding domain**. The key residues in $DT_R$ implicated in HB-EGF binding are highlighted in grey. The corresponding residues in the R domains of CT1 and CT2 are shown in purple and pink, respectively. Residues were aligned structurally in pymol.

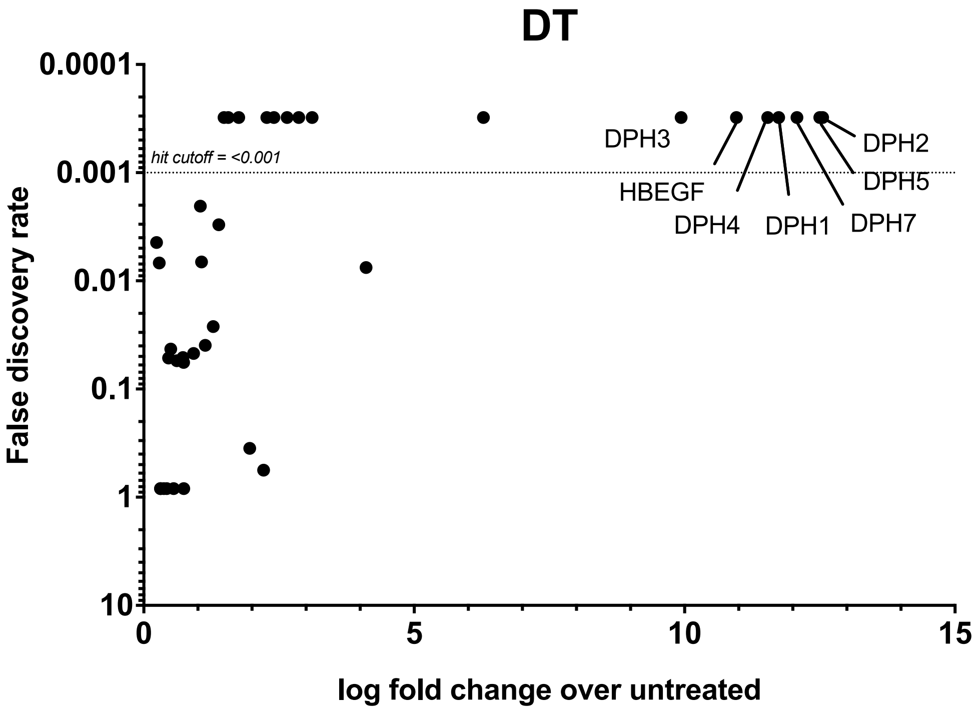


**Supplementary Fig. 5|** CRISPR/Cas9 screen of DT. $EC_{99}$ (10pM) of DT was tested on Hap1 cells, and the top hits were diphthamide synthesis genes, and HBEGF (the receptor).

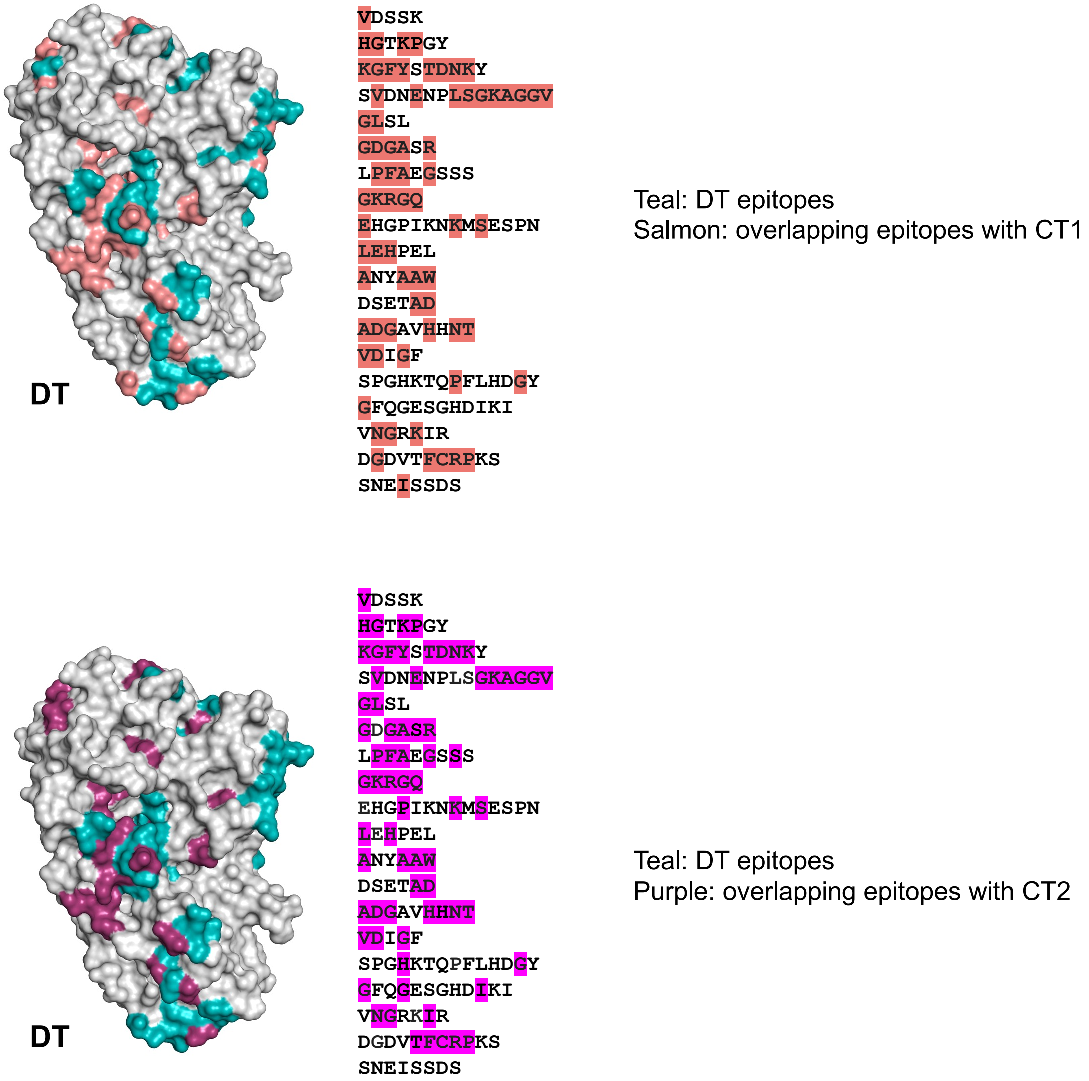


**Supplementary Fig. 6| Epitope residue conservation between DT and CT1 (top) or DT and CT2 (bottom).** The residues implicated in B-cell recognition of DT that are not conserved in CT1 or 2 are highlighted in teal. The residues that overlap with CT1 are highlighted in salmon (top) and the residues that overlap with CT2 are highlighted in purple (bottom). ~50% of the epitope residues are conserved, however only one epitope is entirely conserved in both CT1 and CT2 (GKRGQ).

### Supplemental Figure 7

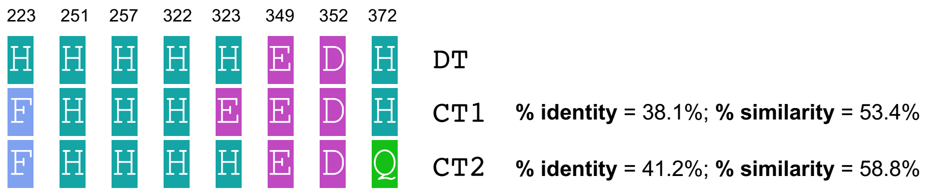


**Supplementary Fig. 7| Sequence alignment of key functional residues in translocation domain**. The key residues in $DT_T$ function are highlighted in grey. The corresponding residues in the T domains of CT1 and CT2 are shown in purple and pink, respectively. Double asterisks represent identical residues between either CT1 or CT2 and DT. Residues were aligned structurally in pymol.
